# Screening of genes related to the interaction between Listeria monocytogenes and host cells

**DOI:** 10.1101/2022.08.28.505572

**Authors:** Gao Fan, Pu Jun-xing, Huang Jin-wen

## Abstract

To investigate how host cells respond to the hijacking of host cells by Listeria monocytogenes (LM) and affect their gene expression in the process of infection by LM. In this study, three data lines in the GEO database were used for differential expression analysis. The results showed that 34 co-expressed genes (DEGs) were selected from the differential expression analysis of three data lines. Among them, 30 genes were co-up-regulated and 4 genes were co-down-regulated. In the blood test group, 131 genes were up-regulated and 28 genes were down-regulated. In the liver, 132 genes were up-regulated and 18 genes were down-regulated. In the spleen, 142 genes were up-regulated and 11 genes were down-regulated. Among the 30 upregulated IDEGs, GBP3, MB21D1, FPR2, SAMHD1, CXCL10, STAT2, IRF1, and STAT1 genes were involved in the defense response to the virus, type i interferon signaling pathway, and inflammatory response. Functional enrichment analysis revealed that 30 up-regulated genes were enriched in cytoplasmic spermatoproteasome complex, proteasome core complex, and cell membrane lateral signaling pathway, which were involved in the regulation of threonine-type endopeptidase activity and guanosine triphosphate (GTP) signaling process.

## Introduction

Listeria monocytogenes (LM) is a foodborne pathogen, which can survive and proliferate in low temperatures, high salt, acidic solution, and other environments. It can cause different degrees of damage to multiple organs or tissues of infected animals, such as blood, liver, spleen, etc., resulting in septicemia, liver necrosis, and other pathological reactions. In addition, LM can also cause a series of clinical symptoms such as abortion and meningitis in pregnant animals [1-2]. In host cells, LM can interfere with or block the normal gene expression of host cells by secreting a variety of toxin factors, to weaken or even lose the immune response to pathogens, and then complete the process of intracellular proliferation and immune escape [3]. In recent years, although LM related pathogenic study more, the research mainly focused on finding new toxin factor, host cell pathological changes, immune escape, etc., and analyses in the infected host cell gene expression differences of related research is relatively scarce, so, in the direction of research and exploration, will help to further understand the relationship between LM and the role of the host cell. In this study, we mined Differential Expression Gene (DEG) in the blood, liver, and spleen of mice infected with LM by using the relevant data of Gene Expression Omnibus (GEO). In addition, the functional enrichment analysis of the screened common Differential Expression genes (IDEGs) was conducted to explore how host cells respond to LM infection by regulating their own Gene Expression. To provide a reference for further study of the mechanism of host cell-LM interaction.

## 1 material

### 1.1 Data and Samples

The data of this study were obtained from the GEO database of the National BioInformation Database of the United States, numbered GSE77102. Female C57BL/6 mice were selected as experimental animals in this data series, and the experimental data of blood, spleen, and liver infected by LM for 1-3D were used as the research objects. The platform used to detect the expression of samples was GPL6887.

### 1.2 Software and Methods

the analysis of the software and the method using GEO database tools GEO2R (https://www.ncbi.nlm.nih.gov/geo/geo2r/), then enter GSE77102, Lm-infected blood test group and normal blood control group, LM-infected liver test group and normal liver control group, LM-infected spleen test group and normal spleen control group were defined. Select False Discovery Rate to correct the P-value. After log2 conversion, differential expression analysis was performed. Transcripts meeting the conditions of “adJ. P < 0.05 and log2FC = ≥1” were set as Differential Expression Transcriptions (DET). According to the information on the GPL6887 platform, the Genes corresponding to DET were found and set as Differential Expression Genes (DEGs). DETs without gene information were deleted. The “biomaRt” package was run with R software (version 4.1.2) to convert the gene name of DEGs. Analysis of LM-infected DEGs in blood, spleen, and liver using the “Veen” package revealed Upregulated, Identical Differential Expression genes in the three tissues. Ur-idea and Downregulated co-differentially expressed Gene (DR-IDEG). The UR-IDEGs and DR-IDEGs were input into the STRING database (http://string-db.org/) for protein interaction analysis, and PPI networks with interaction relationships were obtained. Cytoscape software (version: 3.9.0) was used to visualize the PPI network, and CytoHubba plug-in “Degree” was used to calculate the connectivity between each gene. In this study, the top three genes were selected as the core genes. Use 6.8 (https://david.ncifcrf.gov/tools.jsp) for the UR - IDEGs, DAVID, and DR - IDEGs GO enrichment and KEGG signal pathway analysis, Then using the Sangerbox tool (http://www.sangerbox.com/tool), the analysis results are circle diagram and the bubble chart display.

## 2 Results and analysis

### 2.1 DEGs screened by Data Line In this study

the DEGs of three different tissues of C57BL/6 mice infected by LM were obtained from the GSE77102 data line, as shown in Figure 1. “Log2FC > 1” indicates up-regulated gene expression, while “log2FC < -1” indicates down-regulated gene expression. Table 1 shows the DEGs obtained by screening three different tissues in the data system. 159 DEGs were screened from the blood test group (Figure 1A), among which 131 genes were up-regulated and 28 genes were down-regulated. A total of 150 DEGs were screened from the liver test group (FIG. 1B), among which 132 genes were up-regulated and 18 genes were down-regulated. A total of 153 DEGs were screened from the spleen test group (FIG. 1C), among which 142 genes were up-regulated and 11 genes were down-regulated.

**Figure 1.**
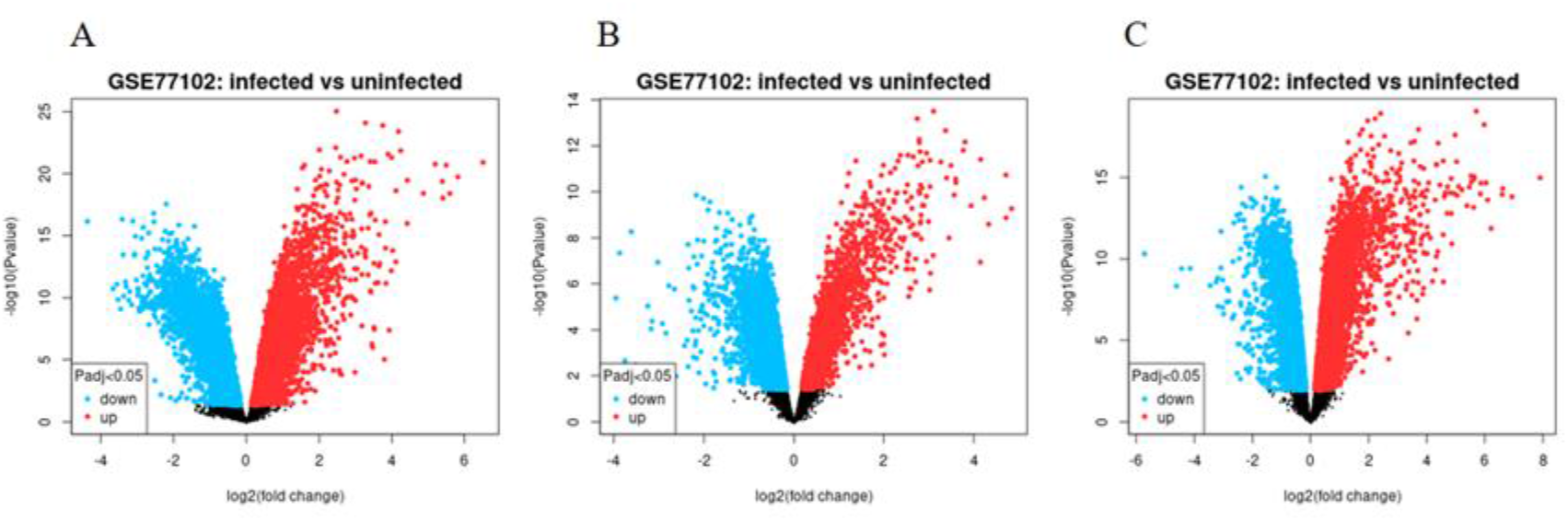
Volcano map of differentially expressed genes in three different tissues.

**Table 1.** Results of differential gene statistics for data lines.

| The data | organi<br>zation | Number of<br>differentially<br>expressed genes / | Number of up-regulated genes / | Number of<br>down-regulated<br>genes / |
| --- | --- | --- | --- | --- |
| GSE7710<br>2 | blood | 159 | 131 | 28 |
|  | liver | 150 | 132 | 18 |
|  | spleen | 153 | 142 | 11 |

### 2.2 IDEGs screened by three kinds of tissues

The IDEGs analysis of three kinds of tissues is shown in Figure 2. A total of 34 IDEGs were screened from the three groups of DEGs (Figure 2A), and 30 UR-IDEGs were screened from the upregulated DEGs (Figure 2B), while no DR-IDEGs were screened from the downregulated DEGs (Figure 2C). In this paper, we focused on these 30 UR-IDEGs. Among these genes with interaction relationships, STAT1, IRF1 and CXCL10 ranked in the top three, which were the core genes screened (FIG. 3).

**Figure 2.**
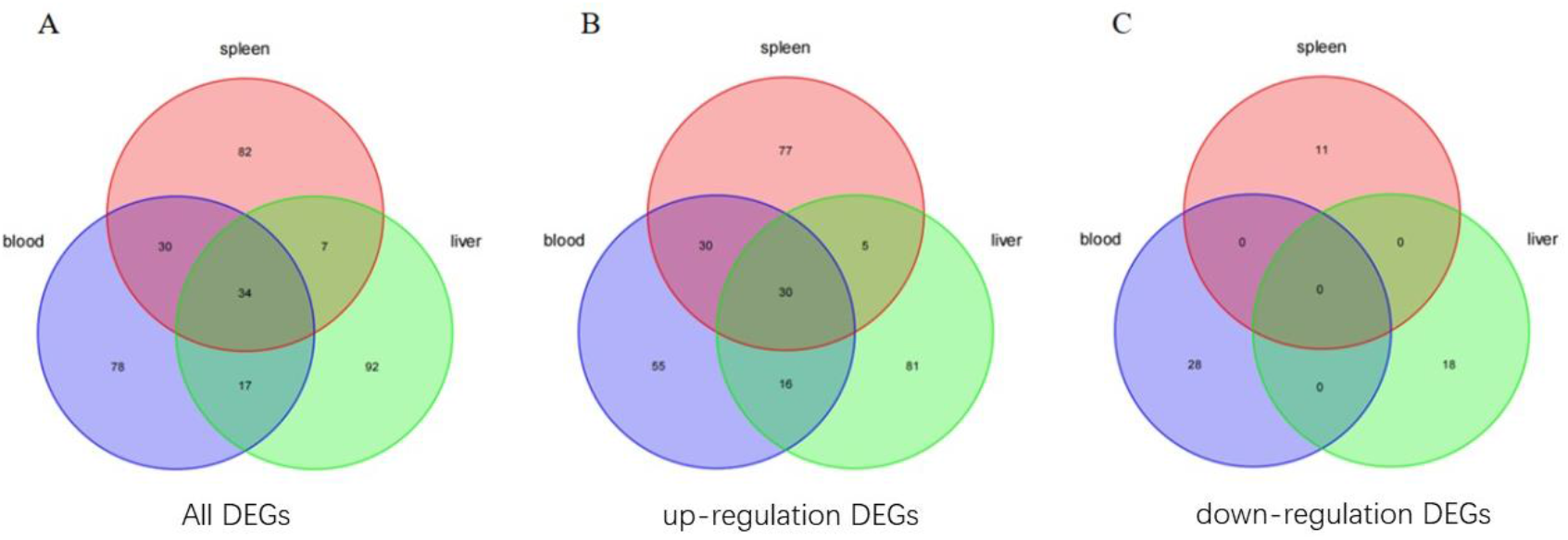
Venn diagram of differentially expressed genes in three experimental groups.

**Table 2.** UR-IDEGS statistics in the three experimental groups.

| Gene name | full name |
| --- | --- |
| ACOD1 | aconitate decarboxylase 1 |
| BATF2 | basic leucine zipper transcription factor, ATF-like 2 |
| CCR5 | chemokine (C-C motif) receptor 5 |
| CD274 | CD274 antigen |
| CXCL10 | chemokine (C-X-C motif) ligand 10 |
| FPR2 | formyl peptide receptor 2 |
| GBP3 | guanylate binding protein 3 |
| GBP7 | guanylate binding protein 7 |
| HDC | histidine decarboxylase |
| HK3 | hexokinase 3 |
| IRGM | interferon gamma-induced GTPase |
| IRF1 | interferon regulatory factor 1 |
| LGALS3 | lectin, galactose binding, soluble 3 |
| MB21D1 | A mab-21 domain containing 1 |
| MLKL | mixed lineage kinase domain-like |
| PARP12 | poly (ADP-ribose) polymerase family, member 12 |
| PHF11 | PHD finger protein 11A |
| PSMB10 | proteasome (prosome, macropain) subunit, beta type 10 |
| PSMB8 | proteasome (prosome, macropain) subunit, beta type 8<br>(large multifunctional peptidase 7) |
| PSMB9 | proteasome (prosome, macropain) subunit, beta type 9<br>(large multifunctional peptidase 2) |
| SAMHD1 | SAM domain and HD domain, 1 |
| SLC15A3 | solute carrier family 15, member 3 |
| SLFN12 | schlafen 1 |
| SNX10 | sorting nexin 10 |
| STAT1 | signal transducer and activator of transcription 1 |
| STAT2 | signal transducer and activator of transcription 2 |
| TAP1 | transporter 1, ATP-binding cassette, sub-family B (MDR/TAP) |
| TAPBPL | TAP binding protein-like |
| UPP1 | uridine phosphorylase 1 |
| WARS | tryptophanyl-tRNA synthetase |

**Figure 3.**
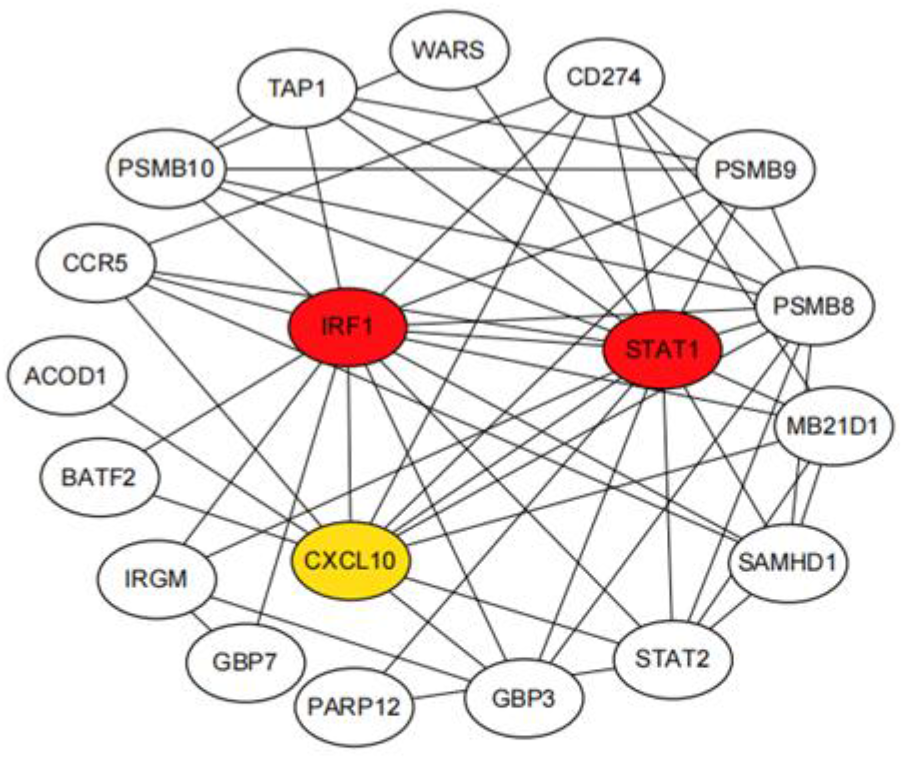
section IDEGs with interaction relationships.

### 2.3 Analysis of functional enrichment pathways of UR-IDEGs Enrichment

analysis was carried out on 30 selected UR-IDEGs, and the condition of screening enrichment pathways was P < 0.05. The results were shown in Figure 4. Among them, the BP approach and KEGG pathway are mainly enriched in defense responses to the virus, beta interferon response, type I interferon signaling pathways, tumor necrosis factor-mediated signaling pathways, involved in cellular protein catabolism process of protein hydrolysis, inflammation, regulating cell amino acid metabolism, antigen processing and through MHC class I, Tap-dependent presentation of exogenous peptide antigens, NIK/NF-κB signaling, proteasome, chemokine signaling, toxoplasmosis, and hepatitis C (Figure 4A). Cellular Component (CC) was significantly associated with the sperpasome complex, the proteasome core complex, the proteasome complex, the cytoplasm, and the lateral side of the plasma membrane, while MF was significantly enriched in THR endopeptidase activity and GTP binding processes (FIG. 4B).

**Figure 4.**
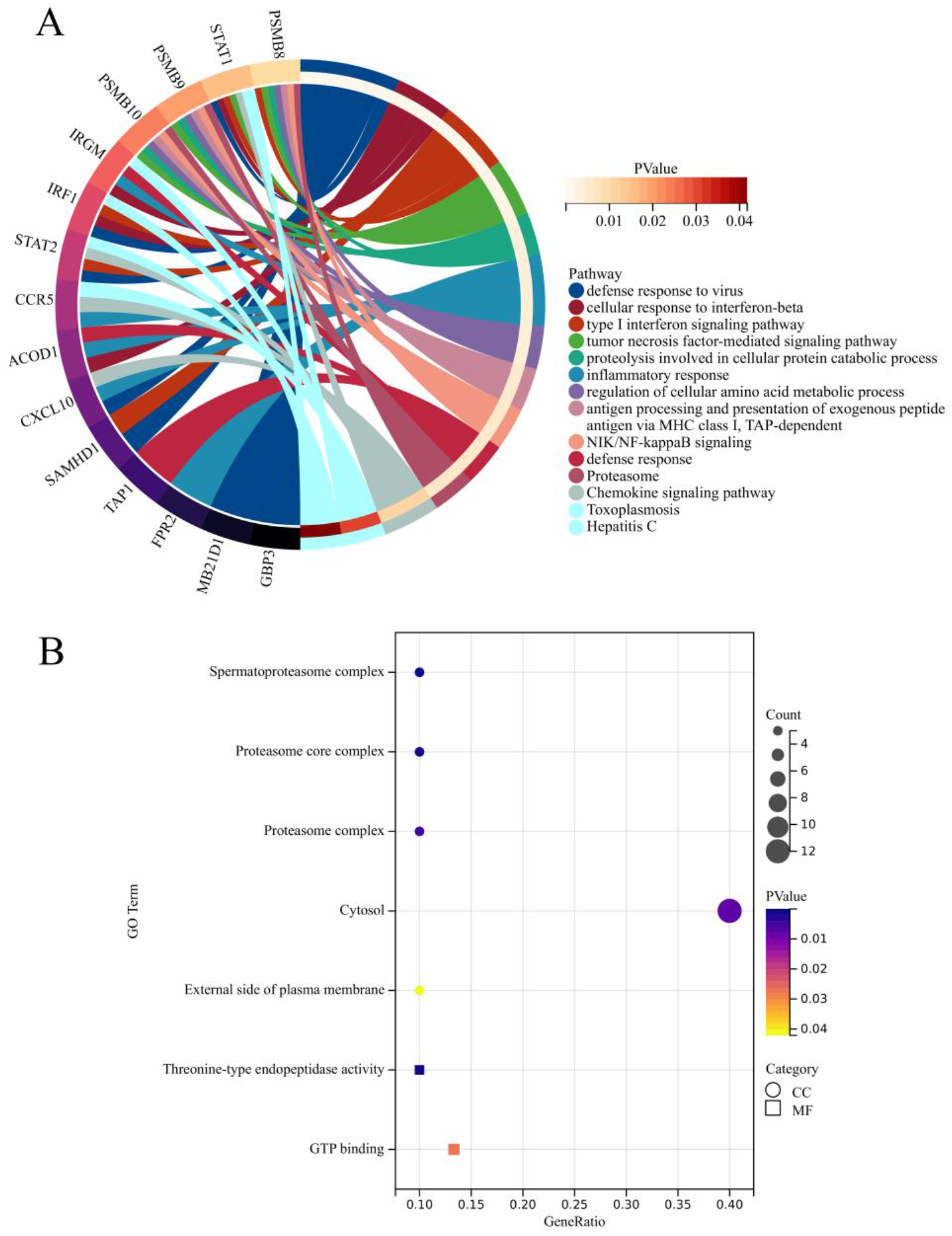
GO and KEGG enrichment analysis results of UR-IDEGS.

## 3 Discussion

### 3.1 Differential expression analysis method and analysis

In this paper, the GEO2R software was used for analysis. The standards and methods of GEO2R software were used for differential expression analysis of the three groups of data, and the transcriptome data information was mined to the maximum extent, to make the co-expressed gene information obtained more reliable and accurate. By differential expression analysis of the transcripts of the three data lines, it was found that the number of up-regulated DEGs was much higher than the number of down-regulated genes in the LM infection group compared with the normal control group. This suggests that there is a complex interaction mechanism between LM and host cells in infected blood, liver, and spleen cells. At the same time, infection will lead to necrosis and apoptosis of more cells.

### 3.2 The regulation strategy of IDEGs involved in the reaction

After LM infects host cells, it will trigger the antiviral response, β-interferon response, and other processes. Wandel M P et al. [4] found that GBP3 plays an important role in maintaining cellular autonomous immunity, and its expression products can regulate the activity of caspase-4, which plays a key role in the defense against bacterial infection. This indicates that GBP3 has a positive role in anti-LM infection. The expression product of MB21D1 is an important effector that extensively acts on the type I interferon response and plays an important role in enhancing the antiviral response of type I interferon [5]. However, in this study, MB21D1 was not directly involved in the process of cell response to LM type I interferon, which may indicate that there are differences in the process of cell anti-LM infection and antiviral infection. Formyl peptide receptor-2 (FPR2) is a seven-transmembrane G-protein coupled receptor that plays an important role in sensing bacterial invasion and modulating immune responses. However, FPR2 is beneficial to the replication of some viruses during infection [6]. In addition, in the process of Streptococcus agalactiae infection, FPR2 can promote the anti-inflammatory effect by regulating the production of chemokines, thereby enhancing the anti-infection ability of host cells [7]. TAP1 participates in and mediates the defense response to LM. In addition, TAP1 plays a certain role in the process of promoting virus replication and infection [8]. Moreover, TAP1 is also believed to play a catalytic role in mediating the occurrence and development of various cancers [9]. This suggests that TAP1 plays a different role in the process of LM and virus infection, and the high expression of TAP1 may accelerate the carcinogenesis process of LM-infected cells, which is helpful to further understanding the carcinogenesis response of LM-infected cells.

SAMHD1 expression products are triphosphate hydrolases, which can degrade intracellular deoxynucleoside triphosphates (dNTPs) to a low level, thereby limiting viral DNA synthesis and further preventing replication of various viruses in host cells [10]. In addition, SAMHD1 also plays a role in repairing DNA damage and alleviating inflammatory responses. It can affect the degradation of initial DNA at the replication fork by regulating the activity of MRE11 exonuclease, and further, activate the DNA replication process. Meanwhile, SAMHD1 can mediate type I interferon response by activating the CGAS-STING pathway [11]. CXCL10 chemokine plays an important role in the process of fighting inflammation and viral infection. It can bind to the receptor CXCR3 and regulate the immune response by activating and recruiting leukocytes (such as T cells, eosinophils, monocytes, etc.) [12]. Studies have shown that aconite acid decarboxylase 1 (ACOD1) plays an important role in immune regulation during inflammation and pathogen infection. ACOD1 is highly expressed in monocytes and macrophages in response to bacterial and viral infections. In addition, ACOD1 has also been confirmed to be highly expressed in LM-infected cells, which is involved in the activation of the β-type interferon response, thereby further resisting LM infection [13]. This suggests that ACOD1 may play an antimicrobial role in the process of LM infection through multiple pathways. CC chemokine receptor 5 (CCR5) is an important G-protein-coupled receptor that can regulate the function of macrophages, T-lymphocytes, and other immune cells and trigger the immune response to pathogens.

In addition, CCR5 is a key receptor for HIV1 and HIV2 to enter host cells [14]. It is not clear whether CCR5 is involved in the process of LM-mediated entry, but the expression of CCR5 is beneficial to enhancing the anti-infection ability of host cells to LM. STAT1 and STAT2 proteins are key mediators of type I and type III interferon signaling, important components of cellular antiviral response and adaptive immunity, and can mediate immune responses through interactions with multiple regulatory factors [15].

Interferon regulatory factor 1 (IRF1) is a multifunctional transcription factor that plays an important role in regulating interferon gene expression and antiviral infection. IRF1 is sharply upregulated after viral infection. It is also involved in mediating type I interferon reaction and type β interferon reaction process [16]. The study of Fehr T et al. [17] found that IRF1 had an anti-infection effect in the mouse model of LM infection, and could mediate the antibacterial response by activating relevant acting factors. IRGM is a negative regulator of the NLRP3 inflammasome, which is beneficial to reduce intracellular inflammation [18]. At the same time, IRGM has also been reported to play a role in the process of Mycobacterium tuberculosis and HIV infection by participating in the process of mediating autophagy to relieve the infection pressure of host cells [19]. PSMB8, PSMB9, and PSMB10 are involved in various responses of host cells in the process of resisting LM infection, indicating that they have an obvious correlation in the synergistic effect. Psmb8-10, both members of the proteasome subunit β family, have been confirmed to play an important role in inhibiting viral replication and promoting viral degradation through the ubiquitin-proteasome system [20]. In addition, the results of enrichment analysis showed that the gene mainly played a role in regulating the activities of various enzymes in the cytoplasm and the signaling of GTP-binding proteins. This indicates that there is a complex interaction mechanism between LM and host cells, and many regulatory processes focus on the response and signal transduction of the host cell membrane to LM infection.

To sum up, the host cell in response to LM infection exists in the process of interaction mechanism is complex, through multiple gene-related functions at the same time also have not been reported before or research, and these commonly expressed genes to the host cell against LM infection plays an important role, and the future further study of those genes could be used to understand the policies of the host cell in LM infections.

## 4 Conclusions

This research use GEO three data in the database system screening LM infection from blood, liver, and spleen cells of differentially expressed genes, screening in 34 common expressed genes, of which are arranged in the top 15 IDEGs main analysis, found its function mainly concentrated in defense responses to the virus, beta interferon response, such as type I interferon response process. Functional enrichment analysis revealed that the genes were mainly enriched in regulating the activities of various enzyme systems and signaling pathways of GTP-binding proteins.

